# Molecular-enriched functional connectivity in the human brain using multiband multi-echo simultaneous ASL/BOLD fMRI

**DOI:** 10.1101/2022.04.21.489018

**Authors:** Ottavia Dipasquale, Alexander Cohen, Daniel Martins, Fernando Zelaya, Federico Turkheimer, Mattia Veronese, Mitul A Mehta, Steven CR Williams, Baolian Yang, Suchandrima Banerjee, Yang Wang

## Abstract

Receptor-Enriched Analysis of functional Connectivity by Targets (REACT) is a novel analytical strategy that enriches functional connectivity (FC) information from functional MRI (fMRI) with molecular information on the neurotransmitter distribution density in the human brain, providing a biological basis to the FC analysis. So far, this integrative approach has been used in blood oxygen level-dependent (BOLD) fMRI studies only, providing new insights into the brain mechanisms underlying specific disorders and its response to pharmacological challenges. In this study, we demonstrate that the application of REACT can be further extended to arterial spin labelling (ASL) fMRI. Some of the advantages of this extension include the combination of neurotransmitter specific information provided by molecular imaging with a quantitative marker of neuronal activity, the suitability of ASL for pharmacological MRI (phMRI) studies assessing drug effects on baseline brain function, and the possibility to acquire images that are not affected by susceptibility artifacts in the regions linked to major neurotransmitter systems.

In this work, we tested the feasibility of applying REACT to resting state ASL fMRI and compared the molecular-enriched FC maps derived from ASL data with those derived from BOLD data. We applied REACT to high-resolution, whole-brain simultaneous ASL/BOLD resting-state fMRI data of 29 healthy subjects and estimated the ASL- and BOLD-based FC maps related to six molecular systems, including the transporters of dopamine, noradrenaline, serotonin and vesicular acetylcholine, and the GABA-A and mGlu5 receptors. We then compared the ASL and BOLD FC maps in terms of spatial similarity, using the Dice Similarity Index and the voxel-wise spatial correlation. On a data subsample (N=19) we also evaluated the test-retest reproducibility of each modality using the regional intraclass correlation coefficient, and compared the two modalities.

Our results showed robust spatial patterns of molecular-enriched functional connectivity for both modalities, moderate to high similarity between BOLD- and ASL-derived FC maps and mixed results in terms of reproducibility (i.e., none of the modalities outperformed the other). Overall, our findings show that the ASL signal is as informative as BOLD in detecting functional circuits associated with specific molecular pathways, and that the two modalities may provide complementary information related to these circuits.

Considering the more direct link of ASL imaging with neuronal acrivity compared to BOLD and its suitability for phMRI studies, this new integrative approach could become a valuable asset in clinical studies investigating functional alterations in patients with brain disorders, or in pharmacological studies investigating the effects of new or existing compounds on the brain.

## 1. INTRODUCTION

Understanding the relationship between the microscale molecular processes of the human brain and its macroscale functional architecture has attracted considerable attention in the neuroimaging community over the years (Cole et al., 2013; Deco et al., 2018; Dipasquale et al., 2019; Hellyer et al., 2017; Rieckmann et al., 2011; Schaefer et al., 2014). The use of multimodal approaches integrating outputs from different techniques and acquisition sequences can enrich the information that can be derived compared to single modalities when studying brain function. This integration has also been fuelled by recent advances in analytical approaches (Deco et al., 2018; Dipasquale et al., 2019) and global data-sharing initiatives (e.g. UK Biobank, ENIGMA Consortium, Human Connectome Project, etc.).

One of the most recent analytical strategies to integrate functional magnetic resonance imaging (fMRI) and molecular imaging is the Receptor-Enriched Analysis of functional connectivity by targets (REACT) (Dipasquale et al., 2019), a novel multimodal method that exploits the temporal dynamics of the fMRI signal to identify functional circuits associated with specific molecular systems. By using neuroimaging templates of receptor density derived from Positron Emission Tomography (PET) or Single-Photon Emission Computerized Tomography (SPECT) imaging in healthy subjects, REACT maps fMRI data onto the space of specific molecular systems and recovers biologically informed networks of functional connectivity (FC).

The application of REACT to clinical and pharmacological blood oxygen level-dependent (BOLD) fMRI studies has demonstrated its ability to uncover new patterns of functional alterations in neurotransmission-related circuits in patients with brain disorders and suggests potential drug-target mechanisms at the system level that could underlie the functional effects of pharmacological treatment (Cercignani et al., 2021; Dipasquale et al., 2020; Dipasquale et al., 2019; Martins et al., 2021; Wong et al., 2021). However, the BOLD signal is an indirect measure of neuronal activity, resulting from the combination of changes in both oxygen consumption and regional cerebral blood flow (CBF), as well as changes in regional blood volume (CBV) (Buxton et al., 2004). Thus, inferring neuronal dynamics directly from the BOLD signal is not straightforward.

Arterial Spin Labelling (ASL) offers an exciting alternative to BOLD since it allows for a more specific measure of global and regional CBF changes associated with neuronal activity (Alsop et al., 2015; Lauritzen, 2001; Wong et al., 1997) and neurotransmitter activity (Dukart et al., 2018; Selvaggi et al., 2019). CBF is also suitable for pharmacological MRI studies assessing drug effects on baseline brain function (Wise and Tracey, 2006), it is acquired with pulse sequences that are resistant to magnetic field inhomogeneity effects (Wang et al., 2004), thus avoiding susceptibility artifacts and signal loss in the orbitofrontal, inferior temporal, and limbic regions that are linked to major neurotransmitter systems (Wang et al., 2011), and its signal is typically well localized to the capillary beds (Liu and Brown, 2007; Ugurbil et al., 2003). However, some of the classical technical limitations of the standard single-shot ASL acquisition techniques (e.g., long tagging time to label inflowing blood and post-labeling delay to allow tagged blood to flow into the brain, which inevitably result in a lower spatial and temporal resolution compared to BOLD imaging) have hampered the use of ASL as a reliable measure of brain FC.

A novel multi-band and multi-echo (MB-ME) ASL/BOLD sequence has been recently developed to overcome some of those limitations and provide high-resolution, wholebrain simultaneous ASL/BOLD data to extend the study of functional connectivity in the human brain (Cohen et al., 2017). This sequence exploits the advantages of MB and ME techniques to improve signal to noise ratio (SNR) and increase the temporal resolution of ASL data. The collection of more than two echoes also allows for the use of advanced data-driven de-noising strategies, including multi-echo independent component analysis (ME-ICA) (Kundu et al., 2013; Kundu et al., 2012), resulting in the identification of reliable resting-state networks, higher CBF/BOLD coupling, and increased signal stability and reproducibility (Cohen et al., 2017; Cohen and Wang, 2019). Data obtained with this MB-ME ASL/BOLD sequence have already been used to estimate and compare ASL- and BOLD-based resting state networks, and to evaluate the coupling of CBF and BOLD signals (Cohen et al., 2017; Cohen and Wang, 2019).

The above mentioned benefits of ASL, its usage as a quantitative surrogate marker of brain function and the latest technical developments of ASL fMRI offer the opportunity to integrate ASL data and molecular imaging with REACT, to gain further insights from underlying neurotransmitter density data and evaluate brain response to pharmacological intervention. If feasible, this application could pave the way to the integration of molecular and functional imaging in pharmacological and clinical studies where resting-state BOLD data were not – or are not intended to be – acquired, providing a biologically-informed tool to complement the evaluation of global and regional changes in CBF.

In this study, we determined the feasibility of using REACT with ASL fMRI data by estimating the ASL-based FC networks related to six molecular systems that have been used in previous BOLD-based REACT studies, and compared the resulting ASL-based molecular-enriched functional networks with those derived from BOLD fMRI data in terms of spatial configuration and test-retest reliability. This novel MB-ME ASL/BOLD sequence that simultaneously acquires ASL and BOLD data offered us the opportunity to perform an unbiased comparison of the two modalities and rule out the possibility that the use of ASL and BOLD data acquired sequentially could lead to the identification of spatial differences in the ASL- and BOLD-derived networks induced by physiological confounds (e.g., different levels of alertness could potentially affect the spatial pattern of the noradrenergic network), not by intrinsic differences between the two fMRI modalities.

## 2. METHODS

### 2.1 Participants and MRI data acquisition

Twenty-nine right-handed healthy volunteers (mean age = 28.0 years, age range 20 – 46, males/females: 9/20) were recruited for this study. Nineteen of them returned at two weeks to repeat the study. Subjects were instructed to refrain from caffeine and tobacco for six hours prior to the MRI exam.

Ethical approval was obtained from the Medical College of Wisconsin/Froedtert Hospital Institutional Review Board, and all subjects provided written informed consent before participating.

Imaging was performed on a 3T MR scanner (Signa Premier, GE Healthcare, Waukesha, WI, United States) with a body transmit coil and a 32-channel NOVA (Nova Medical, Wilmington, MA, United States) receive head coil. Subjects underwent a resting state fMRI scan using a MB-ME PCASL/BOLD EPI sequence (Cohen et al., 2017; Cohen et al., 2018; Cohen and Wang, 2019) with the following acquisition parameters: echo times (TEs) / repetition time (TR) = 11,30,48,67/3500 ms, FOV = 24cm, matrix size = 80×80, slice thickness = 3 mm (3×3×3 mm voxel size), 11 slices with multiband factor = 4 (44 slices in total), FA = 90°, partial Fourier factor = 0.85, and in-plane acceleration (R) = 2, number of time points = 96, total scan duration = 5.6 minutes. PCASL parameters included labelling time = 1450 ms and post labelling delay time (PLD) = 1000 ms. Subjects were instructed to lie still with their eyes closed and stay awake.

A 3D T1-weighted magnetization-prepared rapid acquisition with gradient echo (MPRAGE) anatomical image was also collected to aid with coregistration (TR/TE = 2200/2.8 ms, field of view (FOV) = 24 cm, matrix size = 512 × 512 × 256, slice thickness = 0.5 mm, voxel size = 0.47 × 0.47 × 0.5 mm, and flip angle (FA) = 8°).

### 2.2 Image pre-processing

Data was analyzed using a combination of AFNI (Cox, 1996) and FSL (Jenkinson et al., 2012). First, the anatomical MPRAGE image was coregistered to Montreal Neurological Institute (MNI) space. Each fMRI dataset was then volume registered to the first volume using *mcflirt* in FSL. Importantly, only the first-echo dataset was registered. Subsequent echoes were registered using the transformation matrices from the first echo.

For the BOLD analysis, the four echoes were combined using the 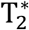-weighted approach(Posse et al., 1999). The data was then denoised using ME-ICA and the open source python script tedana.py version 0.0.10 (https://tedana.readthedocs.io/en/latest)(DuPre et al., 2019; Kundu et al., 2013; Kundu et al., 2012). This technique, described in detail elsewhere, classifies independent components as BOLD or non-BOLD based on whether or not their amplitudes are linearly dependent on TE, respectively (Kundu et al., 2013; Kundu et al., 2012; Olafsson et al., 2015). Non-BOLD components were regressed out of the combined ME data, resulting in a denoised MBME dataset that was then registered to MNI space using the anatomical transformations computed previously. Finally, the data was smoothed using a 6 mm FWHM Gaussian kernel, and high-pass filtered with f_c_ = 0.01 Hz.

For the ASL data analysis, a perfusion-weighted (PW) time series was computed using the first echo by surround subtracting label and control images (Wong et al., 1997).

Finally, global signal regression (GSR) was applied to ASL and BOLD datasets to improve spatial specificity and increase SNR by reducing interindividual physiological differences in ASL data (Jann et al., 2016), and avoid unspecific confounders in BOLD data (e.g., scanner instabilities, physiological drifts, motion, cardiac fluctuations and respiration, level of vigilance) (Birn et al., 2014; Liu et al., 2017; Power et al., 2014; Yan et al., 2013). The global BOLD and CBF signals were calculated using the standard approaches of their respective pipelines, i.e. by estimating for each time point the mean intensity values across the voxels of the whole brain for BOLD data (Fox et al., 2009; Murphy et al., 2009; Power et al., 2014; Zarahn et al., 1997) and grey matter for ASL data (Hodkinson et al., 2014; Martins et al., 2020), given the well-established lack of sensitivity of ASL in the white matter (Hodkinson et al., 2014). We then regressed the global signals out of the BOLD and ASL datasets respectively using *fsl_glm* in FSL.

For completeness, we also performed the subsequent analyses on the BOLD and ASL datasets without GSR and reported the results in the supplementary material.

### 2.3 Molecular-enriched functional connectivity analysis with REACT

Six publicly available in vivo molecular templates of the major neuromodulatory systems, including the transporters of dopamine (DAT) (Garcia-Gomez et al., 2013), noradrenaline (NAT) (Hesse et al., 2017), serotonin (SERT) (Beliveau et al., 2017) and vesicular acetylcholine (VAChT) (Aghourian et al., 2017), and GABA-A and mGlu5 receptors (DuBois et al., 2016; Norgaard et al., 2021) were used as spatial priors to enrich the fMRI analysis of the BOLD and ASL data and estimate the corresponding functional circuits related to each molecular system. We selected these templates to provide illustrative examples of the methodology and its general usage with ASL and BOLD data, although different types of molecular templates can be used in REACT, according to the specific hypotheses of the study and the template availability (e.g., to explore the brain mechanisms underlying a certain disorder, neurotransmitter transporters might be more suited to capture the full architecture of a certain system, while in a drug challenge targeting specific neurotransmitters, the optimal approach would be to use the maps of those specific receptors, if drug binding is known). All molecular templates were previously normalized by scaling the image values between 0 and 1, although preserving the original intensity distribution of the images, and masked using a standard grey matter mask. Of note, the regions used as references for quantification of the molecular data in the kinetic models for the radioligands were masked out of the corresponding template, namely the occipital areas in DAT and NAT and the cerebellum in SERT, VAChT, and mGluR5. The original templates can be found at https://github.com/netneurolab/neuromaps.

A detailed explanation of the REACT methodology and its applications can be found elsewhere (Dipasquale et al., 2019; Martins et al., 2021). In brief, the functional circuits related to the six molecular systems were estimated using a two-step multivariate regression analysis. In the first step, the fMRI volumes were masked using a binarized atlas derived from the molecular data to restrict the analysis to the voxels for which the density information of the neurotransmitters was available in the templates. Then, the molecular templates were used as a set of spatial regressors to weight the fMRI images and estimate the dominant fluctuation related to each molecular system at the subject level. The resulting subject-specific time series were then used as temporal regressors in a second multivariate regression analysis to estimate the subject-specific spatial map associated with each molecular template. The output consists of six maps per subject and fMRI dataset, each one reflecting the molecular-enriched FC associated with a specific neurotransmitter. This analysis was performed using the react-fmri package (https://github.com/ottaviadipasquale/react-fmri) (Dipasquale and Frigo, 2021).

### 2.4 Comparison of BOLD and ASL-based molecular-enriched FC

To estimate population inferences for each neuroreceptor system and fMRI dataset, we performed voxel-wise one-tailed one-sample t-tests using *randomise* in FSL (Winkler et al., 2014) (one test for positive FC and one for negative FC).

The resulting t-stat images of positive and negative FC were compared between datasets using the Dice Similarity Index and the voxel-wise spatial correlation, to assess the similarity between ASL- and BOLD-derived molecular-enriched FC maps in terms of their spatial configuration.

#### Dice Similarity Index (DSI)

After thresholding the t-stat images at p_FWE_ < 0.05 (corrected for multiple comparisons using the threshold-free cluster enhancement (TFCE) option) (Smith and Nichols, 2009), and binarizing them, we measured the degree of overlap between the different datasets using the DSI, calculated as:

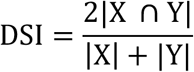

where X are all active voxels in the BOLD-derived FC maps and Y are all active voxels in the ASL-derived FC map. This comparison was run separately for the positive and negative FC (DSI+ and DSI-, respectively).

#### Voxel-wise spatial correlation

The t-stat maps of positive and negative FC were combined in one image per molecular-enriched FC network, down-sampled at 4 mm^3^ to reduce the computational burden of the calculations, and transformed to z-score images. Then, we estimated the correlation of each molecular-enriched FC map obtained from the two datasets using spatial permutation testing (spin test) implemented in the brainSMASH platform (Burt et al., 2020) to account for the inherent spatial autocorrelation of the data.

### 2.5 Test-retest analysis

To quantify the reproducibility of BOLD and ASL response in generating consistent molecular-enriched functional networks, after parcellating each molecular-enriched functional network of all subjects using the Desikan-Killiany atlas and estimating the mean FC of each region, we calculated the regional intraclass correlation between the test and retest sessions for each dataset (BOLD and ASL), ROI and network:

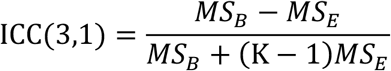

where MS_B_ is the mean square variance between subjects and MS_E_ the mean square error between sessions. MS_B_ and MS_E_ were calculated from an n×k matrix with n=19 observations and k=2 measurements.

In order to compare the regional ICC of the two datasets, the resulting ICC values were Fisher’s z transformed and compared between each possible pair of datasets using paired t-tests (with bootstrapping, 1000 samples). Of note, since the statistical tests were simply applied to help quantify the comparisons between modalities without doing any further statistical inference on the difference of ICC, the results did not need correction for multiple comparisons.

## 3. RESULTS

Figure 1 shows the normalized molecular templates of DAT, NAT, SERT, VAChT, and GABA-A and mGlu5 receptors, and the corresponding significant t-stat BOLD- and ASL-derived FC maps resulting from the one sample t-tests, thresholded at p_FWE_ < 0.05 corrected for multiple comparisons using the TFCE option (Smith and Nichols, 2009).

**Figure 1.**
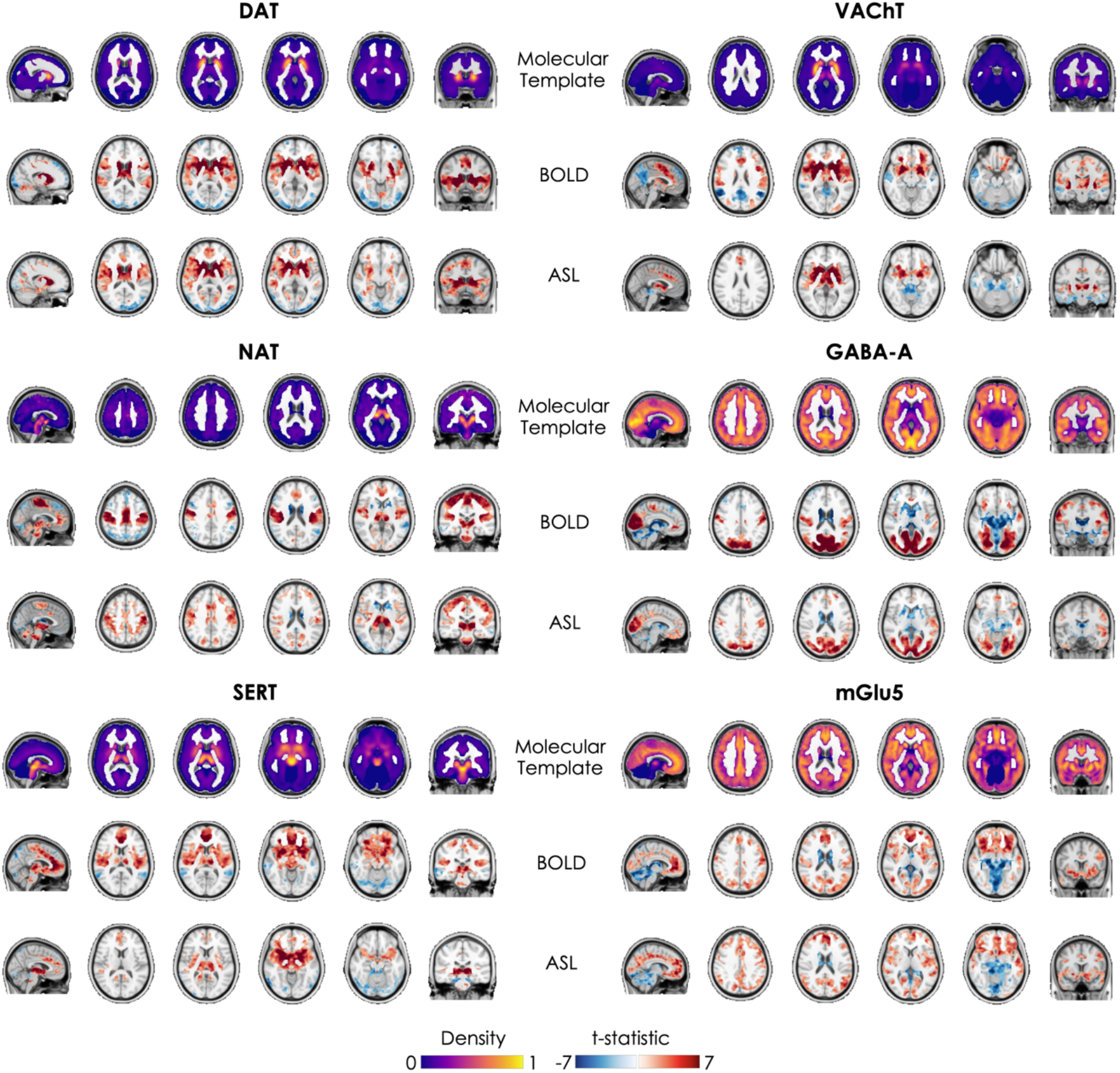
Molecular templates and corresponding molecular-enriched FC maps derived from BOLD and ASL datasets. The distribution density of the molecular templates was rescaled between 0 and 1 and the regions used as references in the kinetic model for all the radioligands were set at 0. The molecular-enriched FC maps are obtained from the significant t-stat maps resulting from the one sample t-tests performed on the positive FC (in red) and on the negative FC (in blue) separately (p_FWE_ < 0.05, corrected for multiple comparisons using the TFCE option).

Overall, the two datasets show good consistency in terms of identification of the main hubs (i.e., regions with positive FC indicating their association with the functional network) of each molecular-enriched network, which align with the distribution of their molecular systems and related pathways. The main hub of the DAT-enriched network corresponds to the striatum (i.e., caudate and putamen), although it is also possible to identify other key cortical and subcortical regions as belonging to this network, including the thalamus, anterior cingulate gyrus, paracingulate gyrus, insula, central opercular cortex, precentral and postcentral gyri, supplementary motor cortex, frontoparietal operculum cortex, superior temporal gyrus and temporal pole. The key regions of the NAT-enriched network identified by both datasets are the brainstem, thalamus, pre- and postcentral gyri, supplementary motor cortex and part of the anterior cingulate cortex, as previously reported in other studies (Cercignani et al., 2021; Martins et al., 2021). The SERT-enriched network is mainly localized in the thalamus, putamen, midbrain, insula, anterior cingulate gyrus and frontal orbital cortex, with the two datasets showing a wider or a smaller involvement of each of these areas. Both datasets identified the striatum, insula and frontal orbital cortex as key regions of the VAChT-enriched network, but significant positive FC was also identified in the pre- and postcentral gyri, supplementary motor cortex and anterior cingulate gyrus. The functional networks related to GABA-A and mGlu5 receptors emerged as more widespread across many cortical and subcortical regions, in line with their wider expression throughout the brain. The areas belonging to the GABA-A-enriched network identified by both datasets are the occipital pole and lateral occipital cortex, lingual gyrus, intracalcarine cortex, cuneus and precuneous, pre- and postcentral gyri, fusiform gyrus and superiol parietal lobule, while the mGlu5-enriched network was mainly localized in the frontal pole, frontal orbital cortex, middle and superior frontal gyri, anterior cingulate gyrus, lateral occipital cortex, occipital pole, paracingulate gyrus, insular cortex, precuneous, occipital pole, angular gyrus, pre- and postcentral gyri and midle temporal gyrus. The complete list of regions having significant within-group positive FC in the molecular-enriched functional maps derived from each dataset can be found in Supplementary Figure 1.

The significant t-stat BOLD- and ASL-derived FC maps resulting from the data without GSR are reported in Supplementary Figure 2. Unlike the networks derived from the BOLD and ASL datasets with GSR, which highlighted, for each molecular system, both the areas of high distribution density of the neurotransmitters and their functional links with brain areas where they project, the areas exhibiting positive FC in the BOLD and ASL datasets without GSR mainly corresponded to those highly enriched with neurotransmitters.

### 3.1 Spatial similarity between datasets

Results from the comparison between datasets estimated using the DSI are reported in Table 1 and visually represented in Figure 2. The BOLD- and ASL-derived maps showed a moderate similarity (DSI range: 0.48 – 0.61) in the spatial distribution of positive FC, and low to high similarity (DSI range: 0.13 – 0.71) in the distribution of negative FC.

**Figure 2.**
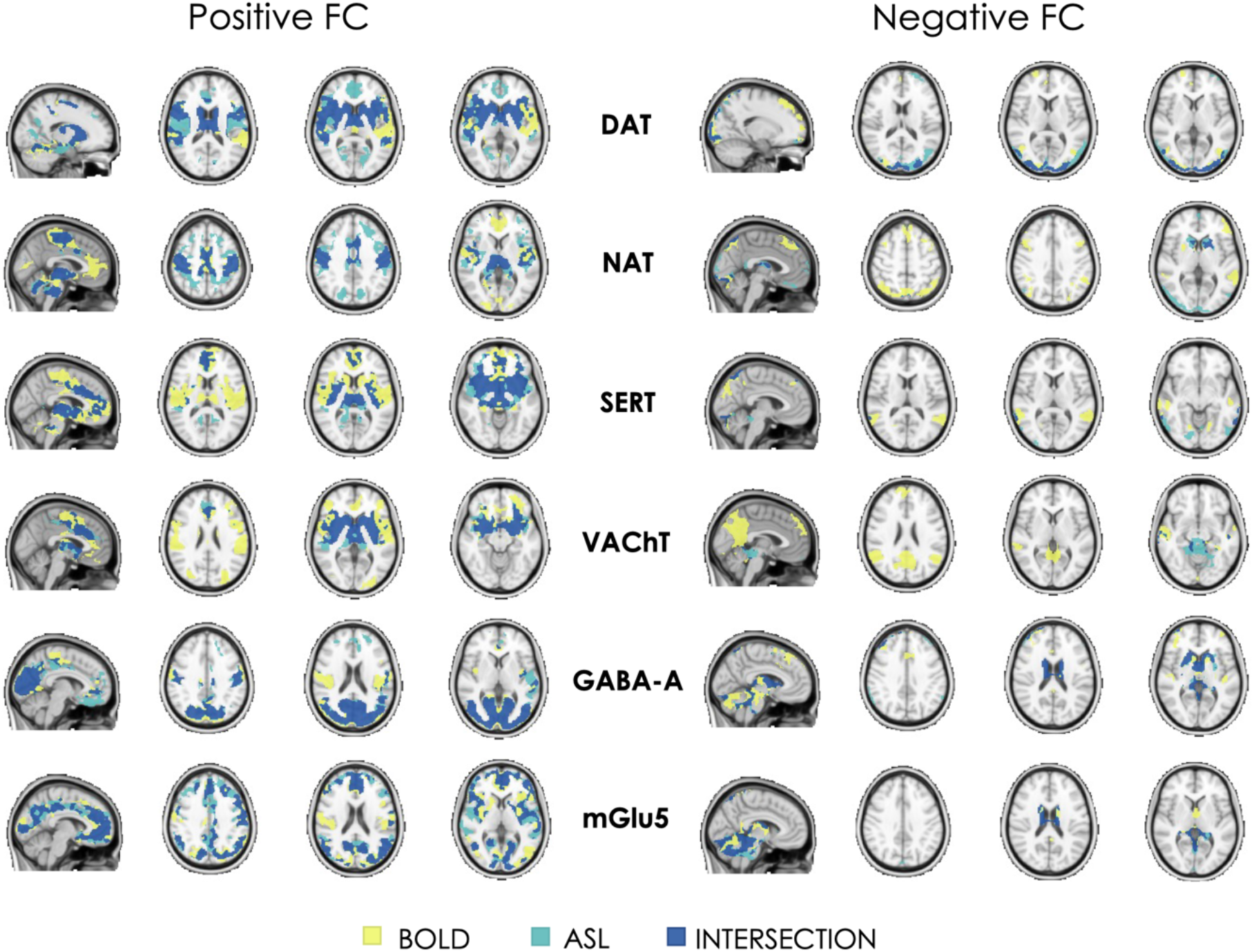
Binary representation of the spatial distribution of the molecular-enriched functional networks derived from BOLD and ASL images and their overlap. The overlap of regions with positive FC and regions with negative FC are represented in two separate panels (on the left and right, respectively).

**Table 1.**
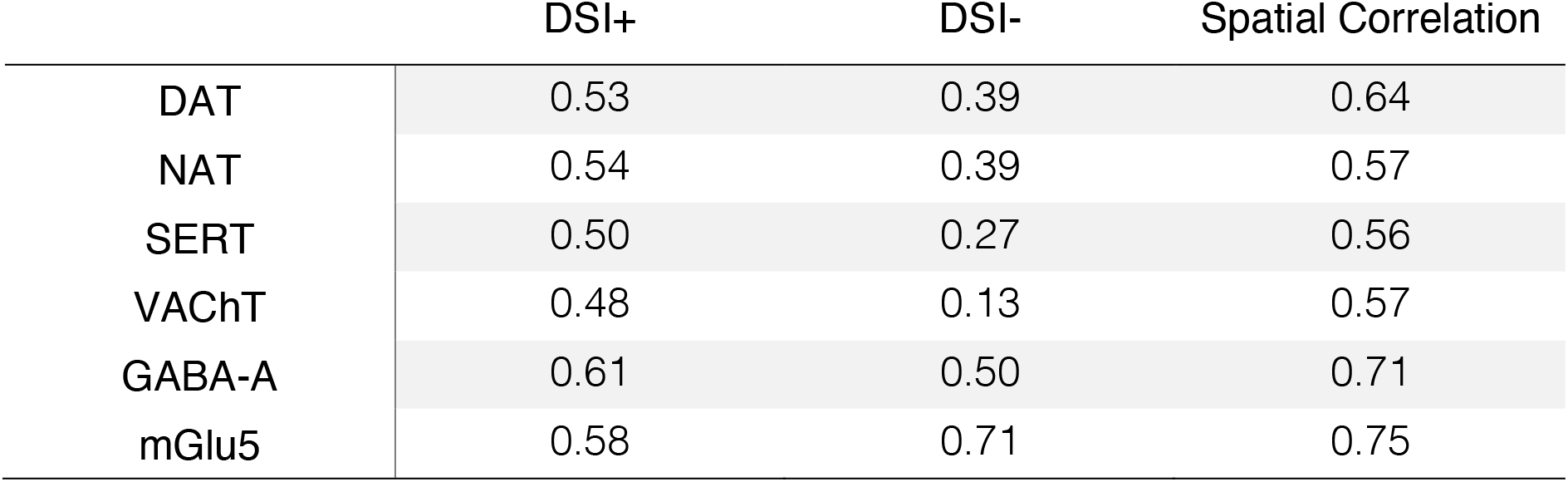
Spatial similarity of the molecular-enriched functional networks between datasets (BOLD and ASL) expressed in terms of Dice Similarity Index (DSI, range: 0-1) and voxel-wise Pearson’s correlation. The DSI was estimated considering the positive (DSI+) and negative FC (DSI-) separately. In terms of voxel-wise Pearson’s correlation, all networks showed significant correlations between datasets (p_spin_<0.001).

In terms of voxel-wise spatial correlation between datasets, we found significant moderate to strong positive correlations between the BOLD- and ASL-derived molecular-enriched FC maps (range of Pearson’s r: 0.56 - 0.75, p_spin_<0.001). All Pearson’s correlations are reported in Table 1.

Of note, the spatial similarity results of the data without GSR are showed in Supplementary Figure 3 and reported in Supplementary Table 1.

### 3.2 Results of the test-retest analysis

Overall, the two datasets showed heterogeneity in the regional distribution of ICC across the whole brain, spanning from very low ICC to moderate ICC (Figure 3). The regional ICCs of the FC maps estimated from the ASL dataset showed significant differences with two out of six FC maps of the BOLD dataset, namely with the VAChT enriched FC maps (ASL > BOLD, t-stat = 2.099, p = 0.039) and the GABA-A-enriched FC maps (ASL < BOLD, t-stat = 3.894, p < 0.001). The comparisons of the other FC maps did not show significant differences (ASL < BOLD contrast, DAT: t-stat = 0.92; NAT: t-stat = 1.122; SERT: t-stat = −0.853; mGlu5: t-stat = 1.199).

**Figure 3.**
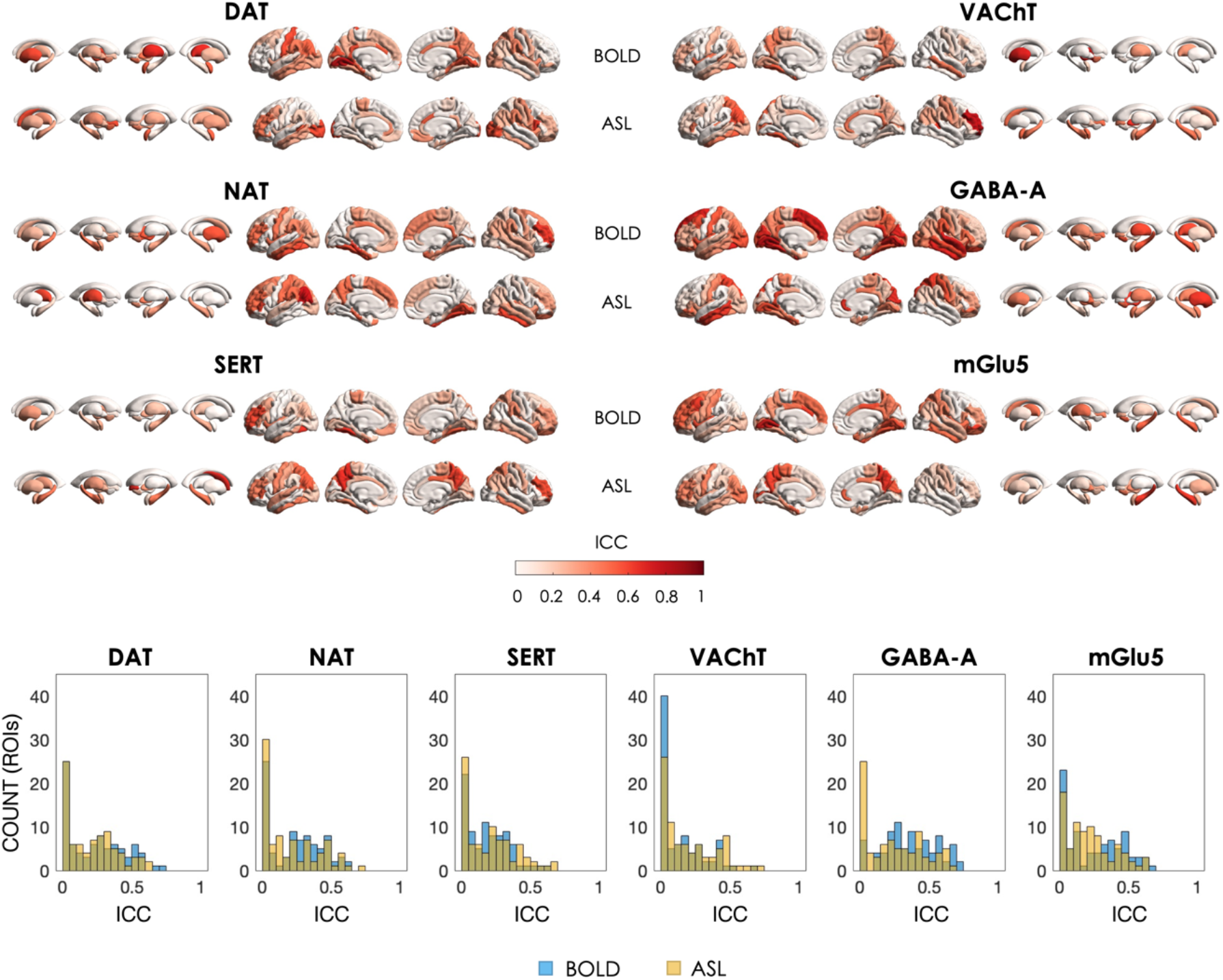
Regional Intraclass correlation (ICC) of the test-retest sessions for each molecular-enriched functional network derived from the BOLD and ASL datasets and corresponding histograms of their distributions.

Of note, the results of the test-retest analysis of the data without GSR are illustrated in Supplementary Figure 4.

## 4. DISCUSSION

In this study, we tested the feasibility of using ASL fMRI data to estimate molecular-enriched functional networks using REACT, a novel analytical method used to project fMRI data onto selected molecular spaces and characterize the functional coactivation of brain regions in terms of their relationship with molecular targets. Specifically, we estimated the functional networks related to six molecular systems (DAT, NAT, SERT, VAChT, GABA-A and mGlu5) using high-resolution, whole-brain resting-state ASL and BOLD fMRI data of healthy subjects acquired with a simultaneous MB-ME ASL/BOLD EPI sequence, and compared the networks resulting from the two modalities in terms of spatial similarity and test-retest reproducibility.

Similarly to the networks derived from BOLD data, the networks resulting from the ASL dataset were able to highlight, for each molecular system, both the areas of high distribution density of the neurotransmitters and their functional links with other brain areas. For example, the main regions belonging to the DAT-enriched functional network estimated from both datasets include the striatal areas, which are the most enriched in dopaminergic innervation, but also specific cortical areas implicated in dopamine modulation of reward, learning, and motivation, such as the anterior cingulate cortex (Arias-Carrion et al., 2010; Ranjbar-Slamloo and Fazlali, 2019), which are functionally and anatomically connected with the striatum (Davey et al., 2012; Di Martino et al., 2008; Haber, 2016). The NAT-enriched functional network shows high positive FC partly in the brainstem, which is the major site of noradrenergic neurons (mainly the locus coeruleus), but also in the thalamus and sensorimotor regions, in line with the modulatory effect of noradrenergic pathways on processing of salient sensory information via actions on sensory, attentional and motor processes (Berridge and Waterhouse, 2003). Importantly, the spatial configuration of such functional networks is also comparable with the functional networks reported in previous REACT-based studies using BOLD fMRI data (Cercignani et al., 2021; Dipasquale et al., 2020; Dipasquale et al., 2019; Martins et al., 2021).

### 4.1 ASL versus BOLD: same but different molecular-enriched networks

Despite the lower SNR of ASL compared to BOLD imaging, our findings show that the spatial patterns of the molecular-enriched functional networks estimated from BOLD and ASL datasets show extensive areas of overlap, as indicated by the moderate network similarity in terms of DSI+ and moderate to high voxel-wise spatial correlation between them. These results confirm the feasibility of using ASL for the molecular-enriched fMRI analysis, but they also show that the networks identified with ASL data do not fully match the BOLD-derived ones. This finding suggests that the two modalities might capture slightly different characteristics of the functional interaction between brain regions within the molecular-enriched networks (e.g. biological differences in the dynamics of spontaneous fluctuations in CBF and BOLD), and could hence be complementary. However, we cannot fully exclude that these differences might also reflect, to some extent, intrinsic differences between modalities, such as SNR or others. For instance, since the BOLD signal originates primarily from the differences in intravascular deoxyhaemoglobin concentration, the spatial correlation to the actual site of neural activity is relatively poor with considerable spatial spreading onto the venous structures (Boxerman et al., 1995; Lai et al., 1993). This phenomenon might, at least theoretically, inflate the amount of shared variance in BOLD signal fluctuations across voxels from the same venous territory. Conversely, ASL typically yields better spatial correlations with the actual site of regional involvement compared to BOLD because the signal originates from smaller calibre vessels (Ugurbil et al., 2003). Further studies will be essential to clarify these between-modality differences and provide additional information on the spatial configuration of the BOLD and ASL networks in other contexts, such as in task fMRI or drug studies.

### 4.2 Reproducibility

In terms of test-retest reproducibility, both datasets showed large variability in regional ICC across brain regions, spanning from low to good ICC, similarly to what has been reported in other fMRI studies (Braun et al., 2012; Cao et al., 2014; Wang et al., 2017). The ASL dataset showed significant differences in ICC with BOLD for two networks, specifically lower ICC in the GABA-A-enriched network and higher ICC in the VAChT-enriched network. This mixed pattern of results does not allow us to provide definitive answers in respect to which modality has a better reproducibility compared to the other modality. However, further studies expanding our approach to task-based fMRI data, where specific brain regions are expected to be engaged, could offer a more comprehensive overview of the reproducibility of these modalities in different contexts.

### 4.3 Limitations

Our study has some limitations worth mentioning. First, the number of subjects was relatively small. However, the purpose of this study was to determine the feasibility of using ASL data to identify molecular-enriched networks comparable to those obtained with BOLD fMRI. Hence, the sample size was appropriate to answer our research question. Second, we explored the spatial configuration of molecular-enriched functional networks of healthy subjects at rest. Although this is the first essential step to define the feasibility of this analysis and pave the way for future works integrating molecular and ASL fMRI imaging, it is important to mention that further studies are needed in order to characterize in detail the similarities and differences between BOLD- and ASL-based molecular-enriched functional networks and investigate whether these might capture distinct aspects of the biology of the brain or simply reflect methodological differences between modalities. This could be tested using pharmacological MRI data to investigate the brain response to a drug acting on selective targets (Handley et al., 2013), or fMRI data with simple tasks that trigger functional changes modulated by specific neurotransmitters (e.g., attentional tasks requiring modulation of the functional network related to the noradrenergic circuit). This would provide better opportunities to explore the specific characteristics of BOLD and ASL functional networks and how they respond to predictable interventions. Ongoing work is already validating this approach in pharmacological MRI, assessing the drug-target mechanisms behind CBF-based functional changes induced by selective serotonin reuptake inhibitors (SSRIs), and in clinical cohorts where a known deficit such as dopamine loss in Parkinson’s Disease likely affects the normal functioning of networks related to specific molecular pathways. Third, our processing pipeline included GSR, although this step is controversial in the fMRI community for both BOLD and ASL imaging and a general consensus has not been reached yet. In ASL imaging, the regression of global CBF is implemented to reduce interindividual physiological differences, which in turn improves spatial specificity and increases SNR (Jann et al., 2016). In the context of BOLD fMRI, the decision to implement GSR is justified by the fact that global fluctuations integrate nuisance components (e.g., scanner instabilities, motion, level of vigilance, cardiac fluctuations and respiration) (Birn et al., 2014; Liu et al., 2017; Power et al., 2014; Yan et al., 2013) that should be removed in order to avoid unspecific confounders - particularly in resting state data and in the context of drug or clinical studies where unspecific vascular or physiological confounders might strongly impact on the results. Other studies have also highlighted that GSR improves the anatomical specificity of FC measures by reducing the contribution of prominent widespread signal deflections in driving high, non-specific inter-regional correlations (Fox et al., 2009), and improves correlations between functional connectivity and behavior (Li et al., 2019). However, it has been shown that GSR also removes – at least in part – neural signals in the data, impacting the configuration of spatially widespread functional networks (Glasser et al., 2016). In the context of multimodal approaches linking molecular and functional imaging, the impact of GSR has not been investigated in depth. We note though that a pharmacological fMRI study exploring the effects of LSD on consciousness showed that GSR significantly improved the spatial match between gene expression maps of LSD targets and functional connectivity changes induced by pharmacological manipulation (Preller et al., 2018). Our results indicate that unlike the networks derived from the BOLD and ASL datasets where GSR was not performed, which showed positive FC mainly in the areas highly enriched with neurotransmitters, the networks resulting from the data with GSR were able to highlight both the areas of high distribution density of the neurotransmitters and their functional links with regions where they project. Overall, although it is not entirely clear why the global signal should affect the identification of links between the molecular substrate of the brain and its functional processes, supported by the pharmacological study mentioned above, our findings suggest that the regression of global signal in BOLD and ASL data is beneficial for a better characterization of the molecular-enriched functional networks. Finally, a relatively short PLD (1.0 s) compared to the recommended PLD of 1.8 s for pCASL (Alsop et al., 2015) was employed for this study. Shorter PLDs can lead to intravascular artifacts if blood does not have adequate time to reach capillary-feeding small arteries. Despite this, this dataset has been used in another study to estimate and compare ASL- and BOLD-based resting state networks (Cohen et al., 2017), showing that resting state networks can be consistently detected using the CBF data. Other studies have also employed short PLDs (Liang et al., 2014) and analyzed FC with a 3D pCASL sequence with PLD = 1.0 s and a BOLD EPI sequence (Jann et al., 2015), finding robust ASL-based connectivity and considerable overlap between ASL and BOLD networks. However, future studies should specifically examine the effect of PLD on resting state FC.

## 5. CONCLUSION

This study shows that the molecular enrichment of ASL fMRI timeseries is feasible and produces molecular-enriched functional connectivity maps that share features with those produced from BOLD fMRI, but also show particularities - which origins we cannot ascertain yet but will need to be explored in future studies. Given the more direct link between ASL perfusion signal and neuronal activity, our integrative approach is likely to become a valuable asset in neuropharmacological studies investigating the effects of new or existent compounds on the brain or in clinical studies investigating functional alterations in brain disorders, particularly when the use of BOLD fMRI might fall short.

## Acknowledgments

MV is supported by MIUR, Italian Ministry for Education, under the initiatives “Departments of Excellence” (Law 232/2016), by Wellcome Trust Digital Award (no. 215747/Z/19/Z). OD, DM, MV, and SW are funded by the National Institute for Health Research Biomedical Research Centre and Clinical Research Facility at South London and Maudsley National Health Service Foundation Trust and King’s College London. The views expressed are those of the authors and not necessarily those of the NHS, the NIHR or the Department of Health.

## Data Availability

Data is available at https://openfmri.org/dataset/ds000216/.

## Code Availability

The react-fmri package can be found at https://github.com/ottaviadipasquale/reactfmri.

