## Supplementary Figure for "Molecular-enriched functional connectivity in the human brain using multiband multi-echo simultaneous ASL/BOLD fMRI"

### Supplementary material

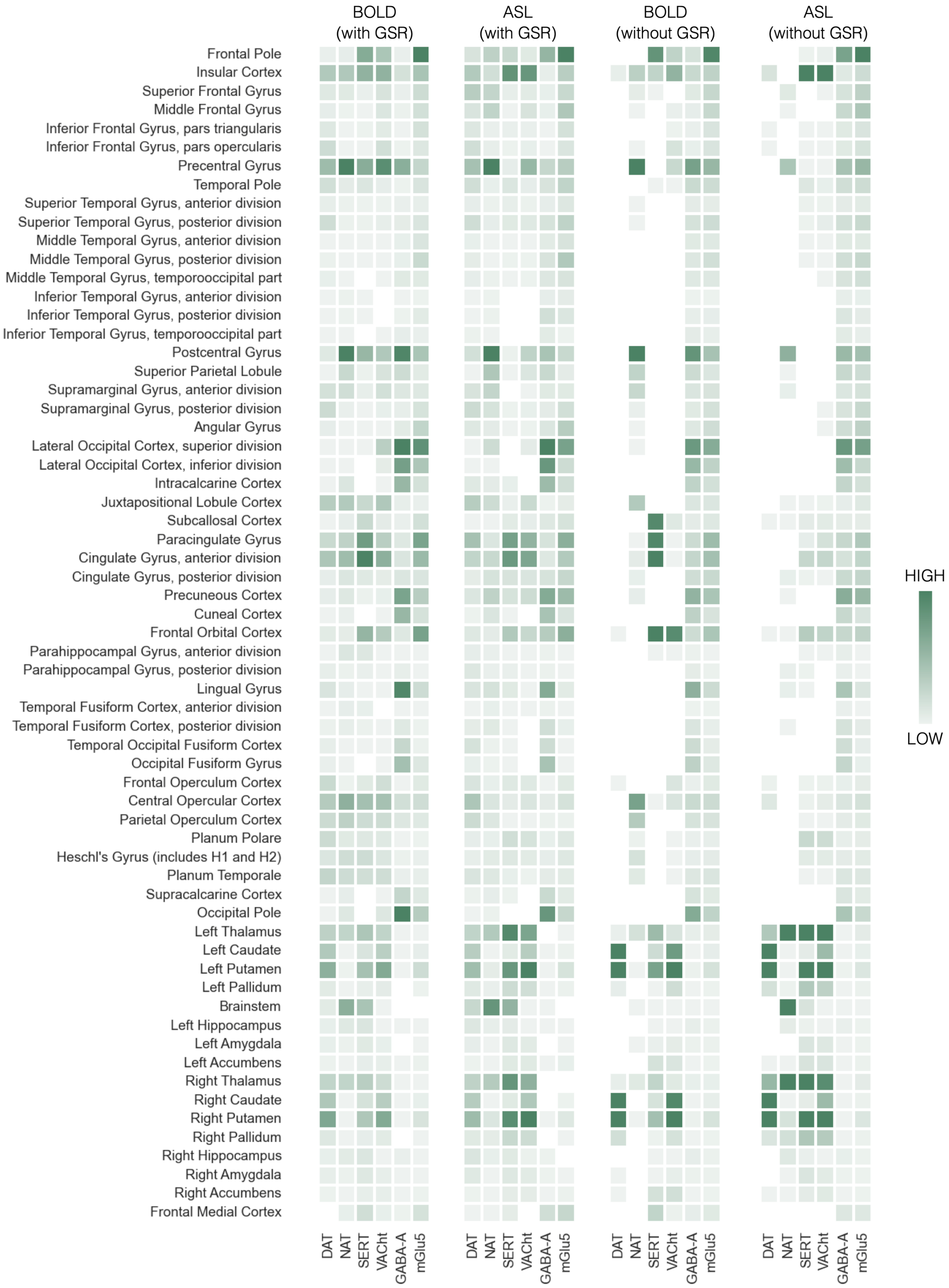


Supplementary Figure 1 **List of regions that reported significant within-group positive FC in the molecular-enriched functional maps derived from the BOLD and ASL datasets (with and without GSR).** For each functional network and dataset, the color scale indicates the probability of the regions to belong to that network.

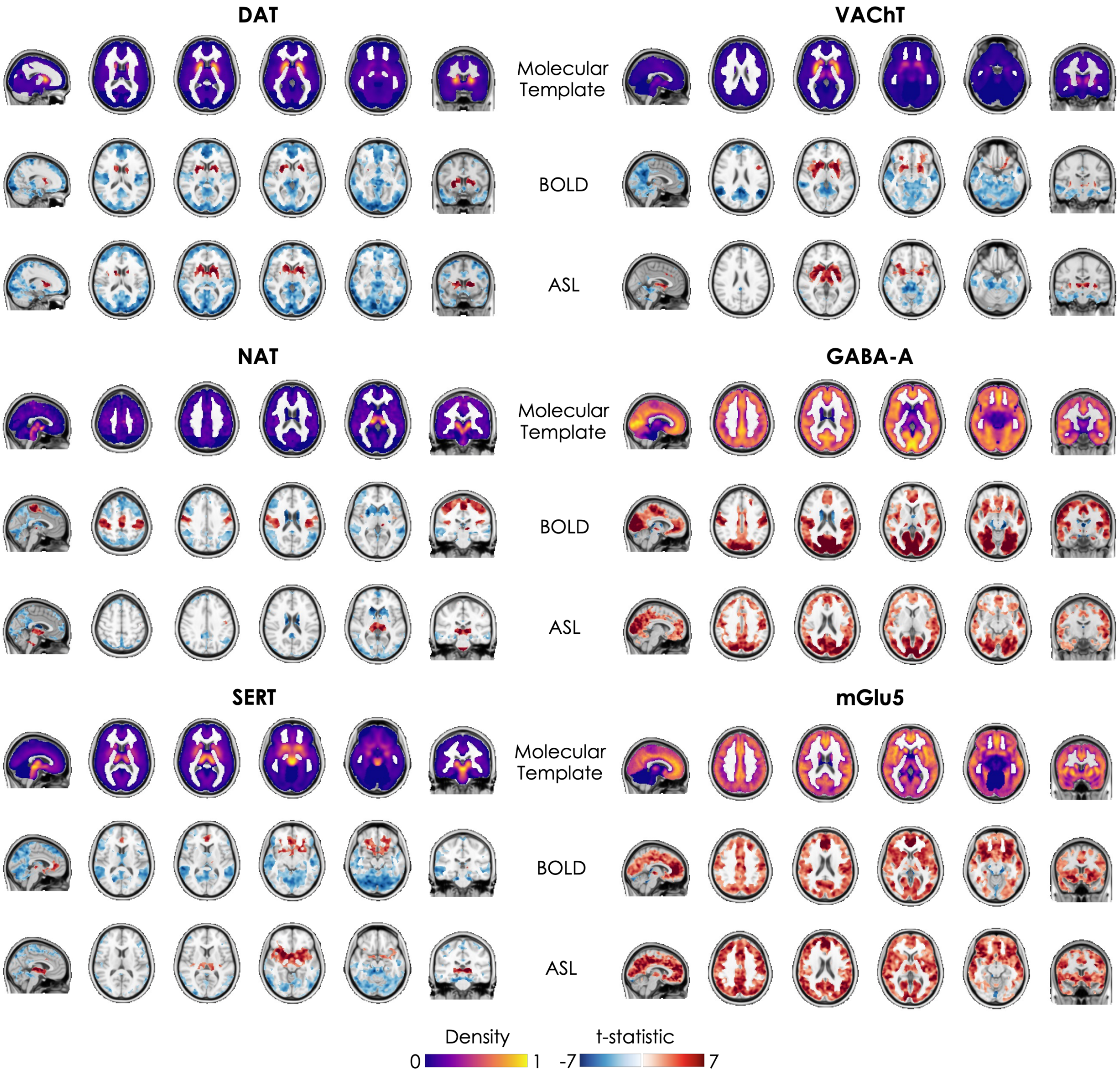


Supplementary Figure 2 **Molecular templates and corresponding molecular-enriched FC maps derived from BOLD and ASL datasets without global signal regression (GSR).** The distribution density of the molecular templates was rescaled between 0 and 1 and the regions used as references in the kinetic model for all the radioligands were set at 0. The molecular-enriched FC maps are obtained from the significant t-stat maps resulting from the one sample t-tests performed on the positive FC (in red) and on the negative FC (in blue) separately ($p_{FWE} < 0.05$, corrected for multiple comparisons using the TFCE option).

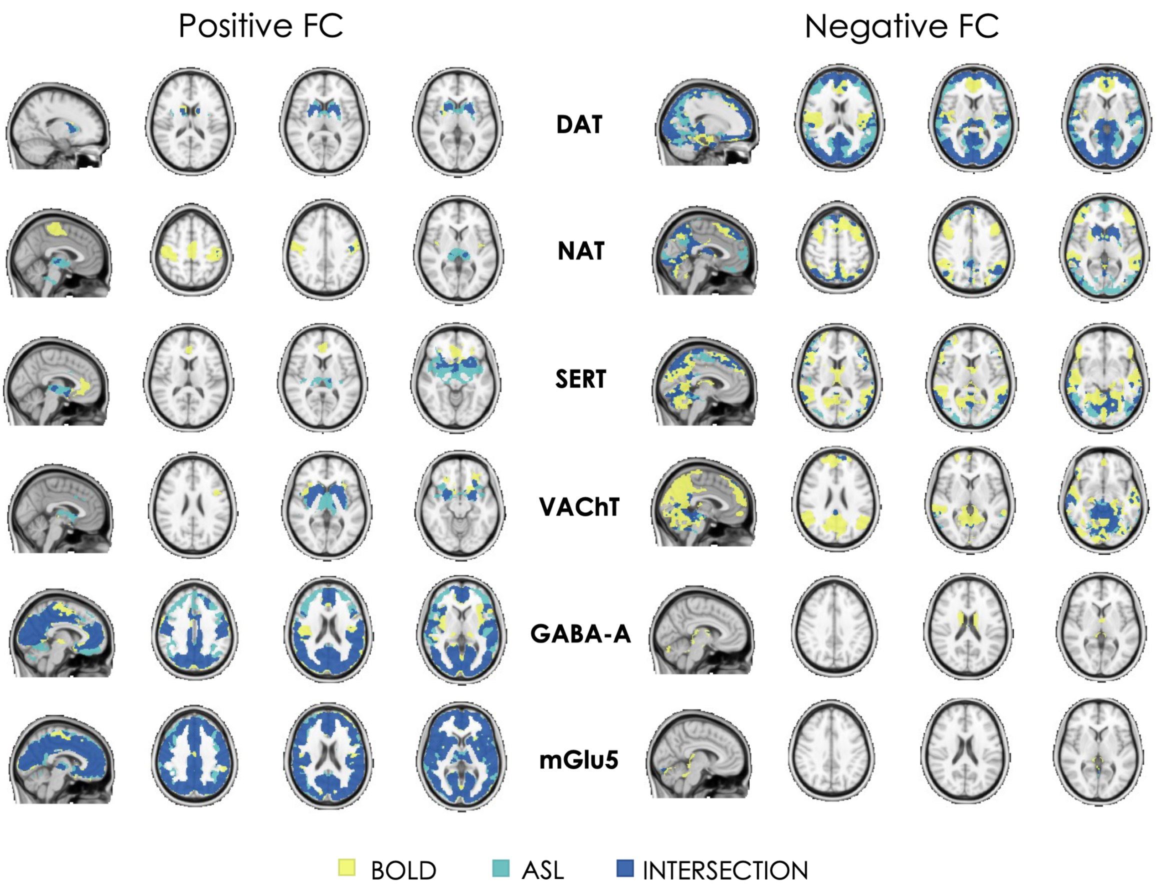


Supplementary Figure 3 **Binary representation of the spatial distribution of the molecular-enriched functional networks derived from BOLD and ASL datasets (without GSR) and their overlap.**

Supplementary Table 1 **Spatial similarity of the molecular-enriched functional networks between BOLD and ASL datasets (without GSR) expressed in terms of Dice Similarity Index (DSI, range: 0-1) and voxel-wise Pearson's correlation.** The DSI was estimated considering the positive (DSI+) and negative FC (DSI-) separately. In terms of voxel-wise Pearson's correlation, all networks showed significant correlations between datasets ($p_{spin}$<0.001).

| | DSI+ | DSI- | Spatial Correlation |
|---|---|---|---|
| DAT | 0.57 | 0.74 | 0.57 |
| NAT | 0.08 | 0.48 | 0.56 |
| SERT | 0.27 | 0.50 | 0.51 |
| VAChT | 0.56 | 0.32 | 0.53 |
| GABA-A | 0.71 | 0.11 | 0.68 |
| mGlu5 | 0.83 | 0.44 | 0.77 |

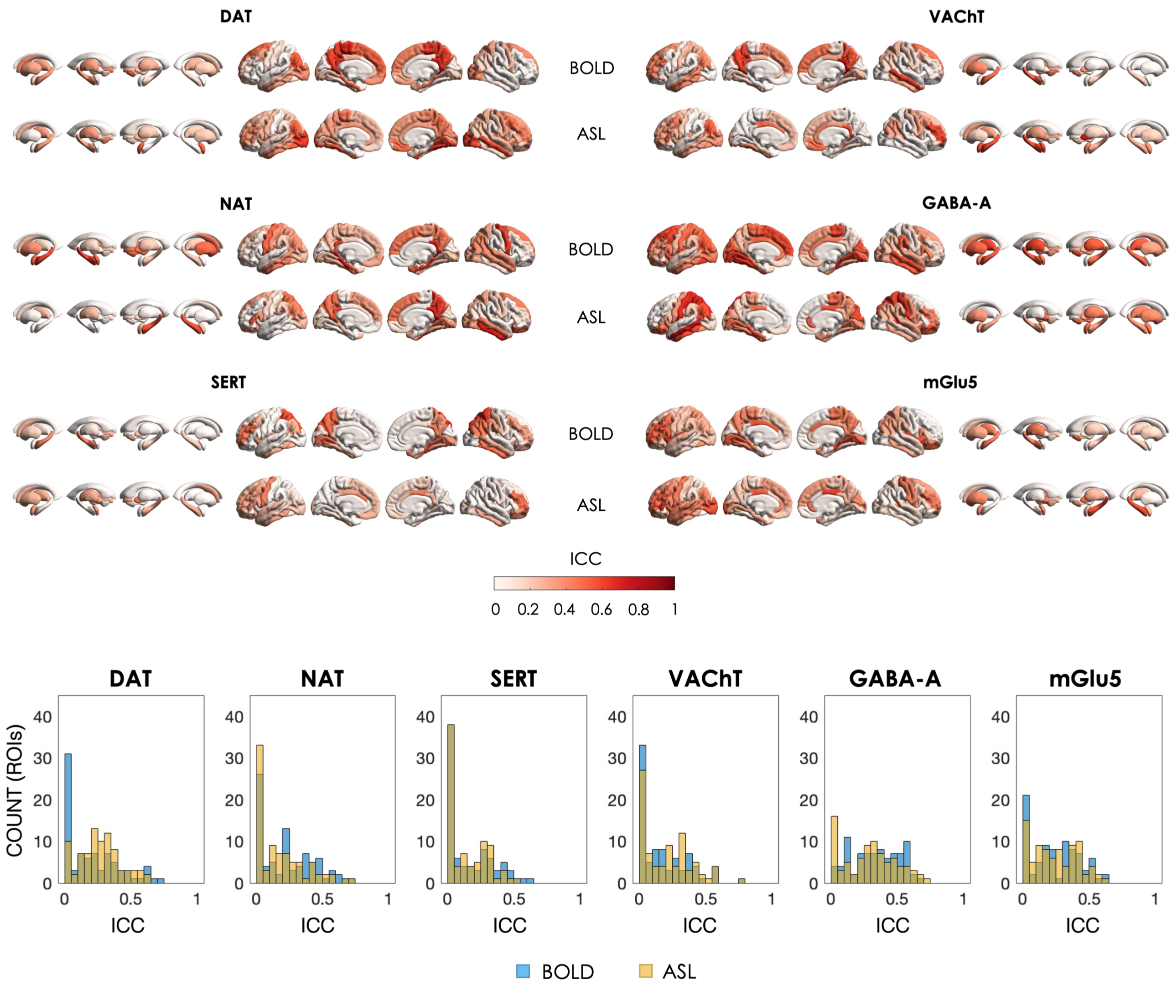


Supplementary Figure 4 **Regional Intraclass correlation (ICC) of the test-retest sessions for each molecular-enriched functional network derived from the BOLD and ASL datasets without GSR and related histograms of their distributions.** The regional ICCs of the FC maps estimated from the BOLD dataset showed significant differences with the BOLD-derived FC maps enriched by DAT (ASL > BOLD, t-stat = 2.119, p = 0.037) and NAT (ASL < BOLD, t-stat = 2.101, p = 0.039).
